# γ-Secretase-mediated endoproteolysis of neuregulin-1 and E-cadherin

**DOI:** 10.1101/2025.04.19.649652

**Authors:** Shweta R. Malvankar, Michael S. Wolfe

## Abstract

γ-Secretase is an intramembrane protease complex, with nearly 150 substrates that are cleaved within their transmembrane domains (TMD). Amyloid Precursor Protein (APP) is the most widely studied, as processive proteolysis by γ-secretase releases the amyloid-β-peptide (Aβ) implicated in the pathogenesis of Alzheimer’s disease. In contrast, proteolysis of other substrates has been little explored. The only known sequence specificity rule for γ-secretase cleavage is for APP, in which phenylalanine is not tolerated at P2’ with respect to any step in processive proteolysis. Recently, we found this specificity rule applies to the initial cleavage of Notch1 substrate as well. In this study, we examined the site of initial cleavage by γ-secretase and explored the phenylalanine rule for two other γ-secretase substrates: neuregulin1 (NRG1) and E-cadherin (CDH1). Upon incubation of recombinant substrates with purified protease complex, followed by mass spectroscopy (MS) and immunoblot analysis, initial cleavage products for NRG1 and CDH1 were identified. Two cleavage sites were observed in the NRG1 TMD, one of which matched that seen previously. However, the observed single CDH1 TMD cleavage site differed from the reported cytosolic cleavage site. Phenylalanine mutants of NRG1 and CDH1 in the P2’ position relative to the first γ-secretase cleavage site showed a shift in the cleavage site, along with reduction in total C-terminal and N-terminal products, compared to that seen with wild-type substrates. Taking together, these findings clarify the initial cleavage sites of NRG1 and CDH1 and support the intolerance of Phe at P2’ position as a general rule for γ-secretase substrates.

**For Table of Contents use only:** 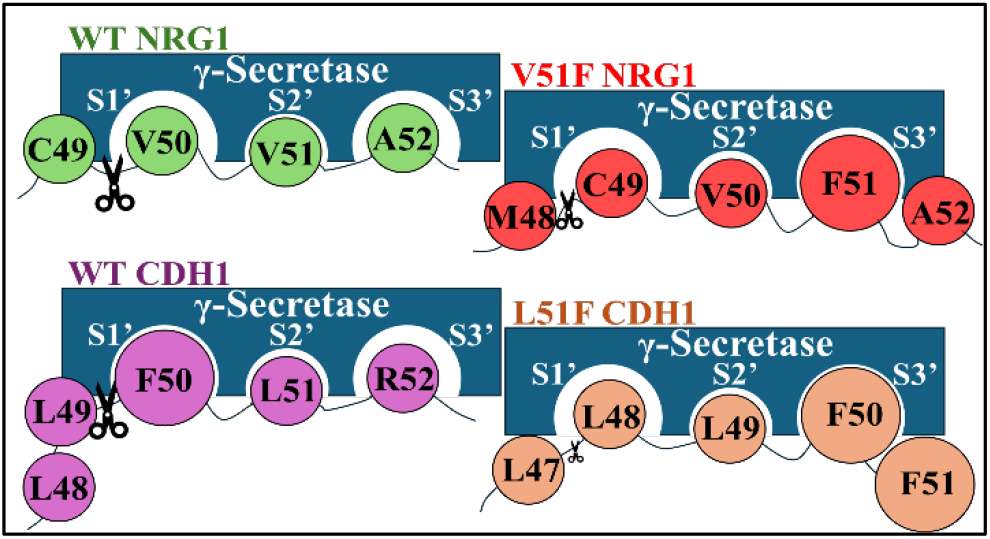

## INTRODUCTION

γ-Secretase is an intramembrane protease complex consisting of four components: Aph1, Nicastrin, Pen2 and the catalytic component Presenilin.^1^ Presenilin contains two aspartates in its active site^2^ that proteolytically cleaves within the transmembrane domain (TMD) of over 145 substrates, including the amyloid precursor protein (APP) and the Notch1 receptor.^3^ APP is cleaved within its TMD multiple times by γ-secretase in roughly tripeptide intervals to release amyloid β-peptides (Aβ),^4,5^ of which the 42-residue variant (Aβ42) aggregates to form cerebral plaques in Alzheimer’s disease. On the other hand, the proteolysis of the Notch1 receptor by γ-secretase is crucial for life. Notch signaling is an evolutionary conserved pathway essential for development and cell-fate determination.^6,7,8^ Aberrant Notch signaling leads to many diseases including cancer.^7,8,9,10^ γ-Secretase substrate neuregulin1 (NRG1) is a ligand for ErbB receptors and plays important roles in development and maintenance of neuronal as well as cardiac systems.^11,12,13^ Disruption of the neuregulin pathway and processing is linked to various neurological and other diseases^13,14^ and is hence considered as a reasonable therapeutic target for the treatment of several disorders.^15,16,17,18^ The intracellular domain (ICD) generated from γ-secretase cleavage of NRG1^19^ likely plays a role in back signaling.^20,21^ Another γ-secretase substrate, E-cadherin (cadherin 1 or CDH1) is essential in maintaining cell-cell adhesion via adherens junctions (AJ). An essential Ca^2+^-dependent component of AJs, CDH1 binds with other components such as catenins to form cell-cell junctions.^22,23,24^ The γ-secretase cleavage of CDH1 is involved in disassembling AJs and releasing the CDH1 ICD,^25^ which can translocate to the nucleus and regulates gene expression.^26^ Dysregulated CDH1 function is associated with various cancers.^27,28,29^

γ-Secretase proteolysis of APP substrate is very well studied, followed by that of Notch1 proteolysis. However, proteolysis of other γ-secretase substrates, even those essential in biology, have not been studied in depth. Fig. 1 depicts the reported first (ε) cleavage sites for APP, Notch, NRG1 and CDH1 by γ-secretase, with corresponding amino acid sequences near these sites. γ-Secretase substrates have no apparent consensus sequence.^30^ The only sequence specificity rule known for γ-secretase cleavage is intolerance of aromatic residues such as phenylalanine at P2’ positions with respect to any cleavage site within the APP TMD.^5^ Very recently, we reported that this ‘phenylalanine rule’ holds true for Notch1 cleavage as well.^31^ However, whether this rule is more general, extending to other γ-secretase substrates is unknown. In addition to the P2’ phenylalanine rule, other common requirements for substrates and their cleavage by γ-secretase includes a short ectodomain, type I intramembrane protein topology, basic and positively charged amino acid residues near the ε-cleavage site (C-terminal end of TMD) and formation of a β-sheet between substrate and PS1 near the ε-cleavage site.^3,32,33^

**Figure 1:**
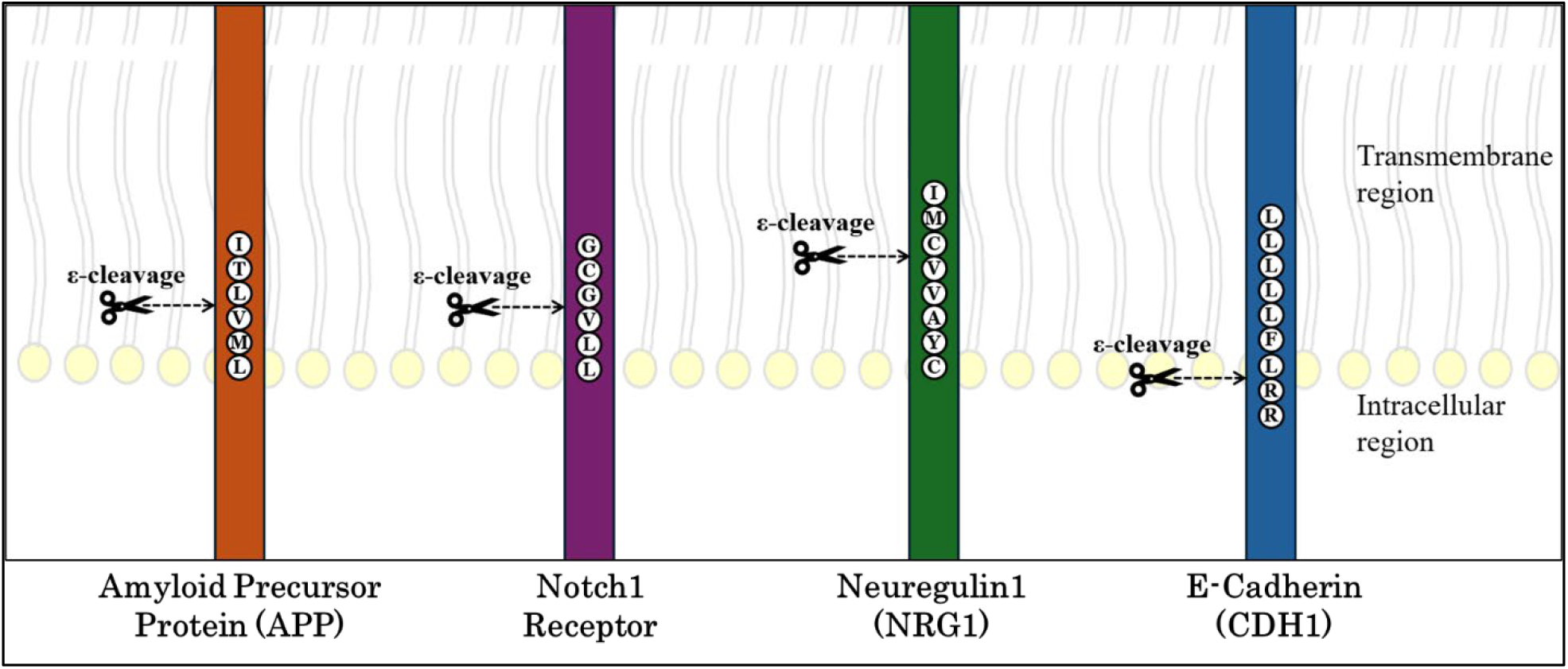
Initial proteolysis of various substrates of γ-secretase. γ-Secretase proteolysis of its substrates: amyloid precursor protein (APP), Notch1 receptor, neuregulin1 (NRG1), and E-cadherin (CDH1) at reported first (ε) cleavage sites are shown. The corresponding amino acid sequences around the cleavage site are shown using single letter codes.

Here we investigated the ε-cleavage sites and the phenylalanine rule for two other γ-secretase substrates, neuregulin1 (NRG1) and E-cadherin (CDH1), via mass spectrometry (MS) and biochemical experiments using enzyme assays with purified proteins. The reported ε-cleavage of NRG1 by γ-secretase occurs well within the TMD,^19^ while that of CDH1 is outside of its TMD (Fig. 1),^25^ making CDH1 an outlier among γ-secretase substrates. We confirmed the reported cleavage site of NRG1 while also identifying an adjacent alternative site. For CDH1, a different cleavage site, within its TMD, was identified, instead of the reported cytosolic cleavage site. We found that phenylalanine is not tolerated at the P2’ position with respect to ε-cleavage for either NRG1 or CDH1. In addition to shifting the initial ε-cleavage site, installation of P2’ Phe reduced formation of cleavage products, although to different degrees for the two substrates. Taken together, these findings identify TMD cleavage sites and validate the only known sequence specificity rule for γ-secretase for two additional biologically important substrates.

## MATERIALS AND METHODS

### Reagents

Anti-flag M2 antibody (Sigma #F1804), HA-tag (6E2) antibody (Cell signaling #2367S), Anti-mouse IgG secondary antibody, HRP (Invitrogen #62-6520)

### Generation of WT and mutant constructs of Neuregulin and E-Cadherin

Both WT and phenylalanine-mutant constructs of neuregulin1 and E-cadherin were designed with an HA tag epitope on N-terminus and a Flag tag epitope on the C-terminus in the pET-21b vector. All the plasmids were obtained from GenScript USA Inc.

### Purification of variants of neuregulin and E-cadherin

*E. coli* BL21 cells were transformed with the respective constructs and plated on LB-ampicillin agar plates and incubated at 37 ^0^C overnight. A single colony was picked for each construct variant and grown with shaking in LB media at 37 ^0^C until OD600 reached 0.8. Cells were induced with 1 mM IPTG and were grown for 3.5 h. Cells were then collected by centrifugation and resuspended in 150 mM NaCl, 10 mM Tris HCl pH 8, 1% Triton X-100. The cell suspension was passed through a French press, and the lysate was incubated with anti-FLAG M2-agarose beads from Sigma-Aldrich (#A2220). The beads were then washed with 150 mM NaCl, 10 mM Tris HCl pH 8, 0.25% NP-40. Bound substrates were then eluted from the beads with 100 mM glycine buffer, pH 2.7, with 0.25% NP-40 detergent and stored at −80 °C. The purified proteins were identified and confirmed using MALDI-TOF MS and SDS-PAGE.

### γ-secretase expression and purification

γ-Secretase was expressed in human HEK Expi293F cells (Gibco #A14635) by transfection with tetracistronic WT pMLINK vector encoding all four components of the protease complex (gift of Y. Shi, Tsinghua University^34^). For transfection, HEK Expi293F cells were grown in suspension in Expi293 Expression medium (Gibco #A1435101) until cell density reached 3 × 10^6^ viable cells/mL. Tetracistronic pMLINK vector (100 µg) was mixed with ExpiFectamine 293 reagent and incubated for 15 min at room temperature and then slowly transferred to the flask with HEK cells. The flask was returned to the incubator-shaker. ExpiFectamine 293 Transfection Enhancer 1 and 2 were added to the flask 21 h post-transfection. The cells were grown and harvested 4 days post-transfection and γ-secretase was purified as described previously.^5^

### Reaction of γ-secretase and substrates

All the enzyme substrate reactions were run in the following manner unless otherwise stated. The purified γ-secretase containing presenilin-1 was incubated for 30 min at 37 °C in the assay buffer containing 50 mM 4-(2-hydroxyethyl)-1-piperazineethanesulfonic acid (HEPES, pH 7.0), 150 mM NaCl, 0.1% 1,2-dioleoyl-*sn*-glycero-3-phosphocholine (DOPC), 0.025% 1,2-dioleoyl-*sn*-glycero-3-phosphoethanolamine (DOPE), and 0.25% zwitterionic detergent 3-[(3-cholamidopropyl)dimethylammonio]-2-hydroxy-1-propanesulfonate (CHAPSO), at final enzyme concentration 30 nM. The substrates were then added to the respective reactions at final concentrations of 10 µM, and the reaction mixtures were incubated at 37 °C for 16 h. The proteolytic products from the enzyme reaction mixtures were then analyzed by mass spectrometry and immunoblotting as described below.

### Detection of C-terminal products by mass spectrometry

C-terminal Flag-tagged ICD products produced from the enzymatic reactions with substrate variants were isolated by immunoprecipitation with anti-FLAG M2 beads (Sigma-Aldrich #A2220) in 10 mM MES (2-(4-morpholino) ethanesulfonic acid) pH 6.5, 10 mM NaCl, 0.05% n-dodecyl-β-D-maltoside (DDM) for 16 h at 4 ^0^C. Flag-tagged C-terminal products were eluted from the anti-Flag beads with acetonitrile: water (1:1) with 0.1% trifluoroacetic acid (TFA). The elutes were run on a Bruker autoflex maX MALDI-TOF mass spectrometer.

### Immunoblotting for detection of intact substrates and products from the enzyme action

Samples from the reactions of γ-secretase and substrates were subjected to SDS-PAGE on bis-tris gels and transferred on PVDF membranes. Membranes were then blocked for 1 h at room temperature in 5% non-fat dry milk in PBST, then incubated with respective primary antibodies for 16 h at 4 ^0^C, followed by washing. The washed membrane was then incubated with corresponding secondary antibodies for 1 h at ambient temperature. Membranes were further washed and imaged for chemiluminescence, and bands were analyzed by densitometry. [Supersignal Western Blot Enhancer (ThermoFisher Scientific #46641) was used to enhance the signal intensity of the N and C-terminal product bands. The PVDF membrane containing product bands were incubated with Antigen Pretreatment solution for 10 minutes followed by washing prior to blocking and primary antibody diluent was used to dilute primary antibody before incubating the membrane in antibody solution.]

## RESULTS

### Design of wild-type NRG1 and CDH1 substrates and their proteolysis by γ-secretase

Based on previous findings that only the TMD of Notch1 or APP substrates is adequate for γ-secretase processing,^35,36,37^ for this study we designed truncated recombinant substrate proteins using human sequences of NRG1 and CDH1. We flanked their TMDs with adjacent short juxtamembrane sequences on each end (Fig. 2A-B), similar to what we reported previously for Notch1 studies.^31^ The protein sequences were further extended with an HA epitope on the N-terminus and a FLAG epitope on the C-terminus to facilitate purification, isolation and identification of the expressed proteins as well as the generated N- and C-terminal products. Using these newly designed constructs, we expressed and purified each recombinant wild-type (WT) protein from BL21 *E*.*coli* bacterial cell suspension by affinity chromatography via the C-terminal FLAG-epitope tag. The purified proteins were then subjected to matrix-assisted laser desorption/ionization time-of-flight mass spectrometry (MALDI-TOF MS) (Fig. 2C-D) and SDS-PAGE gel electrophoresis (data not shown) to confirm their purity and identity.

**Figure 2:**
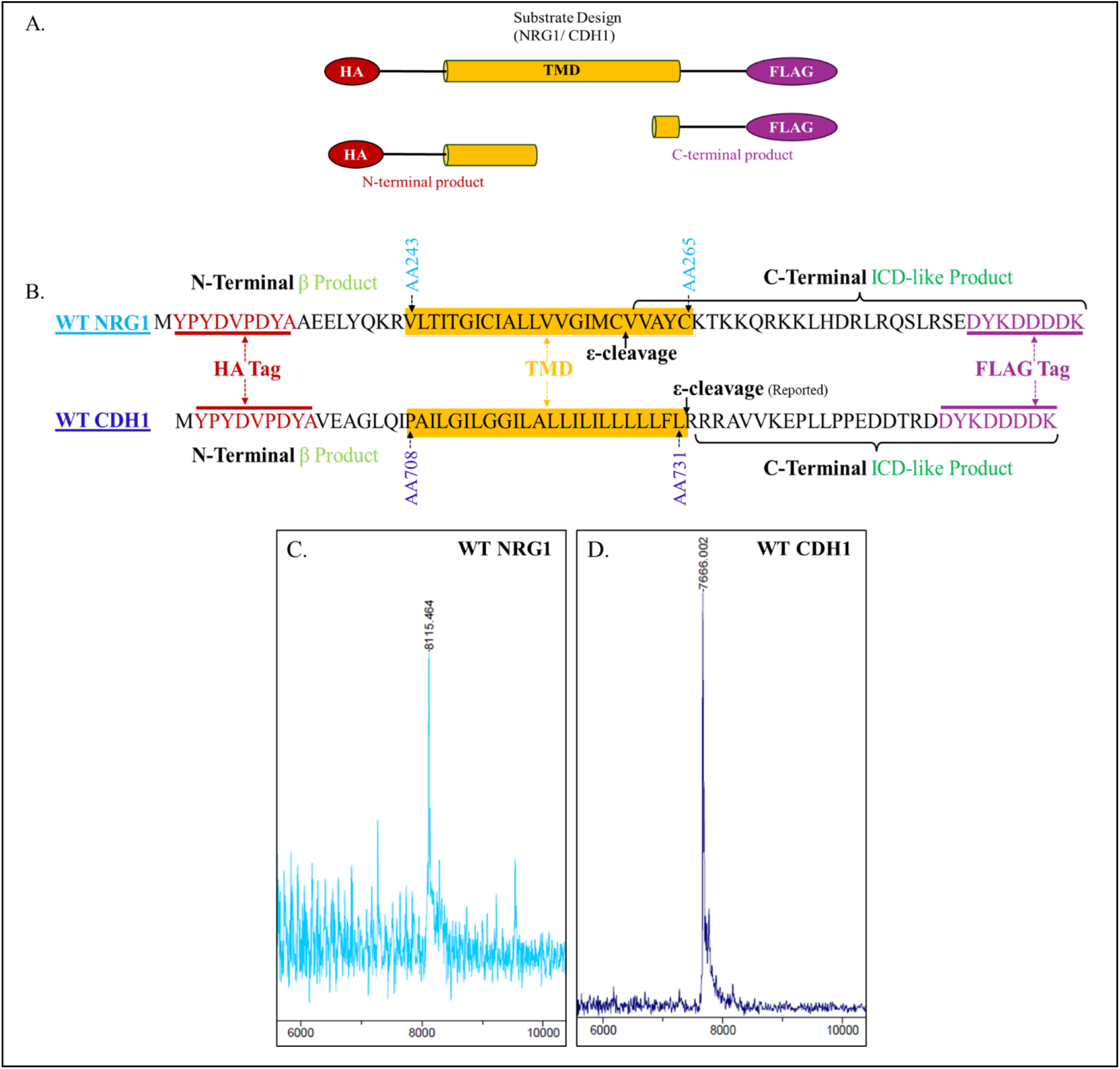
Design and characterization of WT neuregulin1 (WT NRG1) and WT E-cadherin (WT CDH1). A. Schematic representation of truncated WT NRG1 and WT CDH1 substrates tagged on each end (N-terminal HA tag and C-terminal Flag tag), with corresponding C- and N-terminal products. B. TMD sequences for each WT substrate with juxtamembrane sequences on N- and C-termini as well as HA and Flag tags used in this study. The reported γ-secretase ε-cleavage sites for WT NRG1 and WT CDH1 are also shown. The TMDs of NRG1 and CDH1 are shown with original amino acid numbering per their UniProt entries; for NRG1 (TMD: AA243-AA265) and CDH1 (TMD: AA708-AA731), which corresponds to (TMD: AA32-AA54) and (TMD: AA28-AA51), respectively, for NRG1 and CDH1 according to the numbering based on APP cleavage site (shown later in Table 1). MALDI-TOF MS spectra for C. intact purified WT NRG1 substrate and D. intact purified WT CDH1 substrate confirm protein identities. For intact WT NRG1, calculated mass: 8114.450, observed mass: 8115.464; for intact WT CDH1, calculated mass: 7664.981, observed mass: 7666.002.

To verify normal γ-secretase processing of the purified truncated WT NRG1 and WT CDH1 substrates, both substrates were incubated with recombinant WT γ-secretase enzyme that had been expressed in human embryonic kidney (HEK) Expi293F suspension cells and purified as previously described.^5,38^ 30 nM WT enzyme was incubated with 10 µM of the respective WT substrate for 16 h at 37 ^0^C. The formation of C-terminal products (intracellular domain, ICD) was then confirmed by western blot using anti-FLAG antibody for each substrate (Fig. 3 A-B). In both immunoblots product formation was observed in enzyme-substrate reaction mixtures, but not in the presence of γ-secretase inhibitor LY411,575^39,40^ or in control reactions without enzyme or without substrate, confirming that product formation is strictly dependent on enzyme action.

**Table 1.** Alignment of TMD sequences of NRG-1 and CDH-1 with amyloid precursor protein (APP) and Notch-1. TMD sequences were aligned based on observed initial γ-secretase cleavage sites for each substrate and numbered based on APP substrate C99.

| Numbering | 29 | 30 | 31 | 32 | 33 | 34 | 35 | 36 | 37 | 38 | 39 | 40 | 41 | 42 | 43 | 44 | 45 | 46 | 47 | 48 | 49 | 50 | 51 | 52 |
| --- | --- | --- | --- | --- | --- | --- | --- | --- | --- | --- | --- | --- | --- | --- | --- | --- | --- | --- | --- | --- | --- | --- | --- | --- |
| **APP** | G | A | I | I | G | L | M | V | G | G | V | V | I | A | T | V | I | V | I | T | L | V | M | L |
| **Notch 1** | Q | L | H | F | M | Y | V | A | A | A | A | F | V | L | L | F | F | V | G | C | G | V | L | L |
| **NRG1** | Q | K | R | V | L | T | I | T | G | I | C | I | A | L | L | V | V | G | I | M | C | V | V | A |
| **CDH1** | A | I | L | G | I | L | G | G | I | L | A | L | L | I | L | I | L | L | L | L | L | F | L | R |

**Figure 3:**
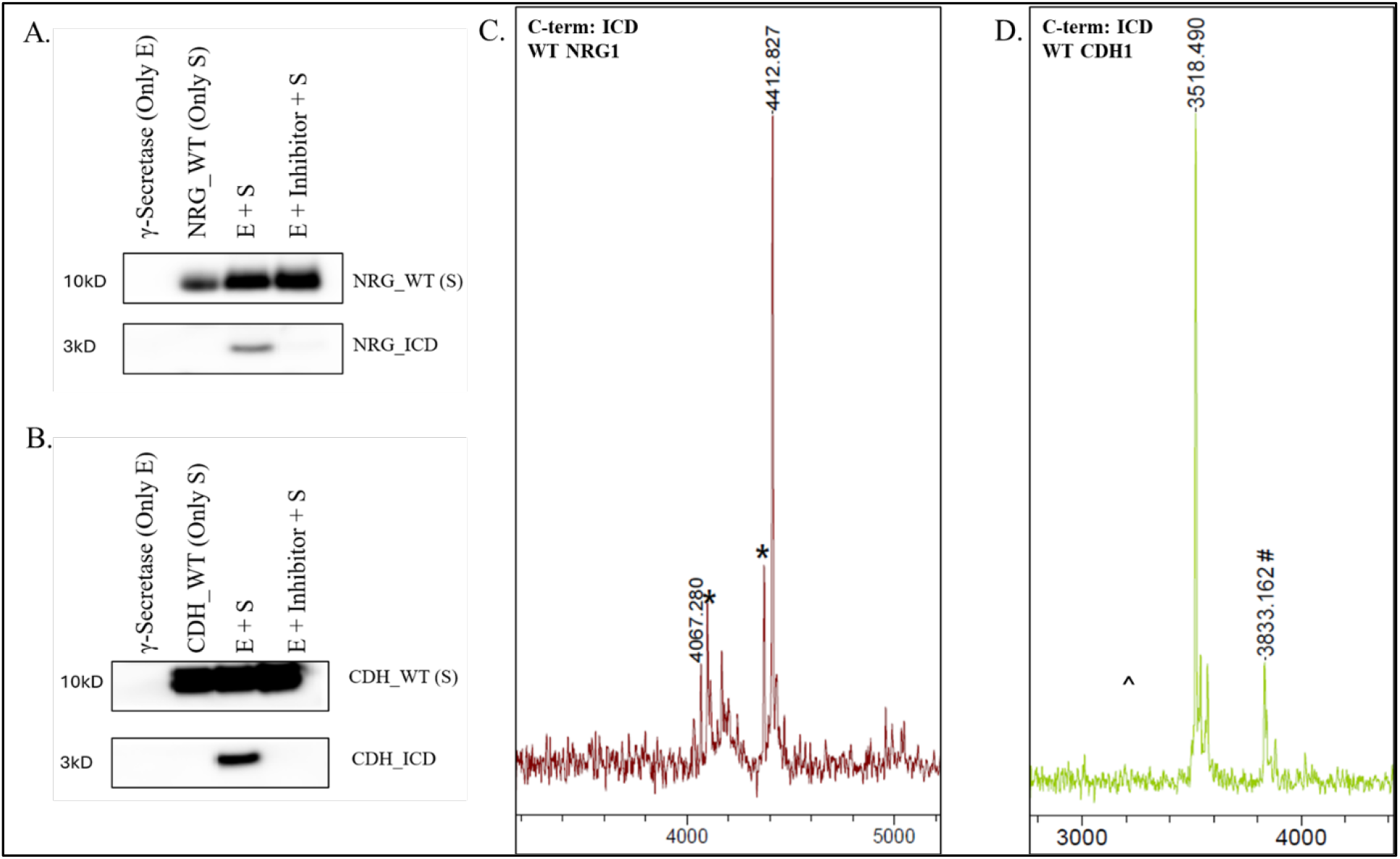
Immunoblots and MALDI-TOF MS spectra for intracellular domain (ICD) formed from γ-secretase proteolysis of WT NRG1 and WT CDH1. Immunoblot analysis of γ-secretase processing of (A) WT NRG1 and (B) WT CDH1 using anti-Flag antibody to detect ICD-like products. In both A and B, no product formation was observed in control reactions with enzyme (E) or substrate (S) alone or in presence of γ-secretase inhibitor (I). C. MALDI-TOF MS spectrum of immunoprecipitated FLAG-tagged C-terminal products (ICDs) from γ-secretase cleavage of WT NRG1. Two cleavage products were observed: (VVAYCKTKKQRKKLHDRLRQSLRSEDYKDDDDK), calculated mass: 4064.123, observed mass: 4067.280; and (IMCVVAYCKTKKQRKKLHDRLRQSLRSEDYKDDDDK), calculated mass: 4411.257, observed mass: 4412.827. D. MALDI-TOF MS spectrum of immunoprecipitated FLAG-tagged C-terminal product (ICD) from γ-secretase cleavage of WT CDH1 showed only one major peak for product (FLRRRAVVKEPLLPPEDDTRDDYKDDDDK), calculated mass: 3515.753, observed mass:3518.490. # represents a peak corresponding to (M+2H)^2+^ of the intact uncleaved WT CDH1 substrate. *Unidentified peaks. In WT CDH1 spectra, ^ represents that no product corresponding to previously reported cleavage site (RRRAVVKEPLLPPEDDTRDDYKDDDDK calculated mass: 3255.601) was observed for WT CDH1.

Further, to detect and confirm the known initial cleavage sites for NRG1 and CDH1, we analyzed the C-terminal products formed from the reactions by MALDI-TOF MS. Immunoprecipitation of the C-terminal products from the enzyme-substrate reactions was performed using beads with immobilized anti-Flag antibody. Based on the C-terminal ε-cleavage products observed for WT NRG1 and WT CDH1, we aligned the corresponding sequences of WT NRG1 and WT CDH1 with the initial cleavage sites of APP and Notch1 previously described (Table 1).^31^ All the amino acid residues of the substrates in this report refer to this numbering system.

The MALDI-TOF spectrum of the NRG1 ICD-like products showed cleavage of the substrate TMD at two distinct sites (Fig. 3C). One cleavage site was observed at the previously reported site between C49-V50,^19^ with another cleavage site between G46-I47, three amino acids in the N-terminal direction. Cleavage between G46-I47 has not been reported before, perhaps because our proteolytic reactions were performed with purified enzyme and substrate. Interestingly, the MALDI-TOF spectrum of CDH1 ICD-like products did not show cleavage at the previously reported site, L51-R52.^25^ In this previous report, Edman degradation was used to determine the sequence of the ICD generated from cultured cells. We suggest that the γ-secretase-generated ICD may have been subsequently trimmed on its N-terminus by an intracellular protease or trimmed after cell lysis by an adventitious protease. Instead, ε-cleavage occurred between L49-F50 (Fig. 3D), within the TMD of the CDH1 substrate. This observation is consistent with the membrane-embedded catalytic aspartates of γ-secretase and cleavage of other substrates within their TMDs.

### Phenylalanine is not tolerated at substrate P2’ position

After ε-cleavage analysis of the WT substrates and confirming the initial ε-cleavage sites for NRG1 and CDH1, we designed their phenylalanine mutants to test the ‘phenylalanine specificity rule'. Phenylalanine was placed in each WT substrate in the P2’ position relative to the first ε-cleavage observed (that is most C-terminal), to generate V51F NRG1 and L51F CDH1 (Table 2). Expression and purification followed by confirmation of these Phe-mutant proteins were performed as noted earlier for WT proteins (Fig. S1).

**Table 2.**
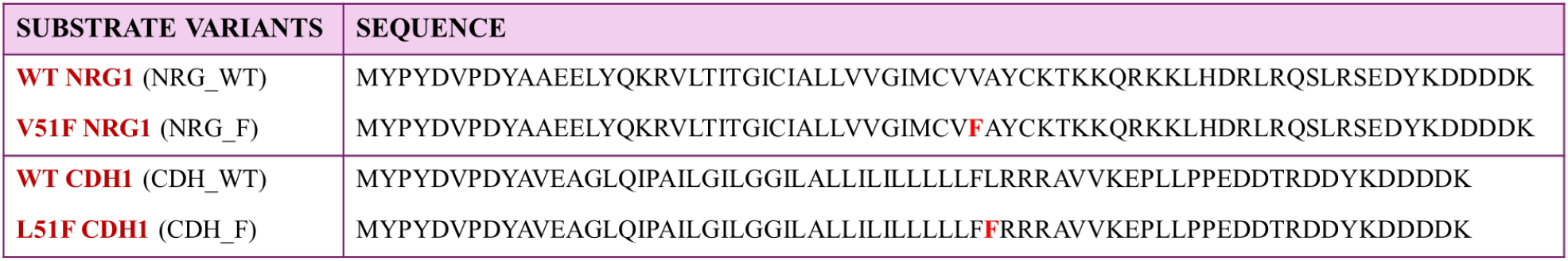
Sequences of NRG1 and CDH1 and their Phe mutants used in this study.

These mutant proteins were then subjected to reaction with γ-secretase and immunoprecipitated using anti-FLAG antibody as mentioned earlier for WT proteins. The MALDI-TOF spectrum of V51F NRG1 ICD-like products showed a clear shift in the cleavage from C49-V50 to M48-C49 (Fig. 4A). No product was seen for C49-V50 cleavage, hence avoiding Phe in the P2’ position. However, the cleavage between G46-I47 was not affected by Phe at 51 position, as Phe does not coincide with P2’ position relative to this cleavage site (Fig. 4A). All these observations are consistent with the Phe specificity rule.

**Figure 4:**
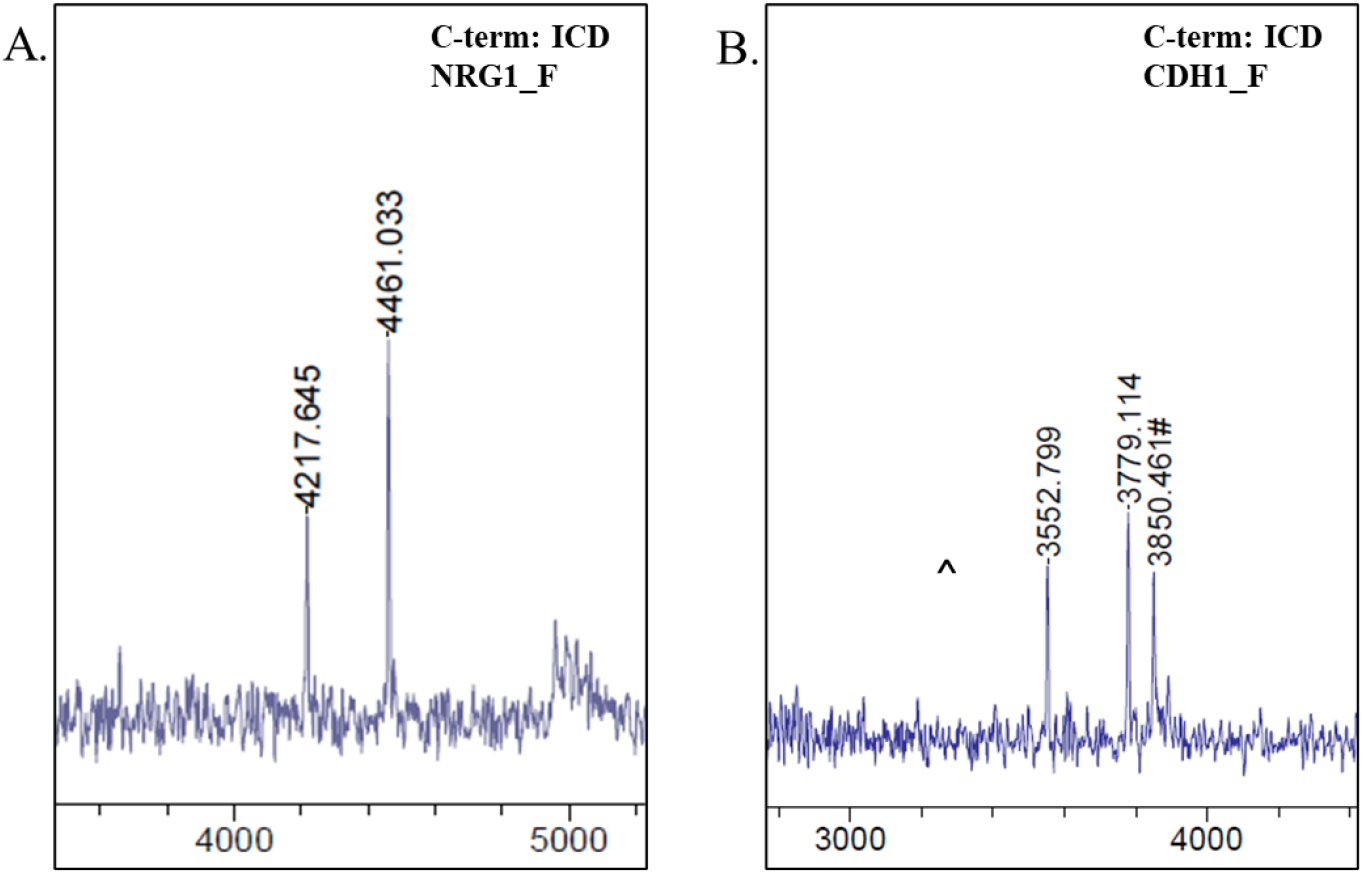
MALDI-TOF MS for phenylalanine mutants of NRG1 and CDH1. A. MALDI-TOF MS spectra of immunoprecipitated FLAG-tagged C-terminal products (ICDs) from enzyme reaction with V51F NRG1 substrate. Two small peaks were observed: for CVFAYCKTKKQRKKLHDRLRQSLRSEDYKDDDDK, calculated mass: 4215.132, observed mass: 4217.645; and for IMCVFAYCKTKKQRKKLHDRLRQSLRSEDYKDDDDK, calculated mass: 4459.257, observed mass: 4461.003. B. MALDI-TOF MS spectra of immunoprecipitated FLAG-tagged C-terminal products (ICDs) from enzyme reaction with L51F CDH substrate. Two small peaks were observed: for LLFFRRRAVVKEPLLPPEDDTRDDYKDDDDK, calculated mass: 3775.906, observed mass: 3779.114; for FFRRRAVVKEPLLPPEDDTRDDYKDDDDK, calculated mass: 3549.738, observed mass:3552.799. The peak for FRRRAVVKEPLLPPEDDTRDDYKDDDDK, calc mass: 3402.669 was not observed. #Observed peaks correspond to (M+2H)^2+^ of the respective uncleaved substrate. ^Product that would have resulted from cleavage at the previously reported site, RRRAVVKEPLLPPEDDTRDDYKDDDDK, calculated mass: 3255.601, was not observed in the spectra of L51F CDH1.

The MALDI-TOF spectrum of ICD-like cleavage products from the enzyme reaction with L51F CDH1 showed a shift in the cleavage site exactly by two amino acids to the left instead of one, showing cleavage at L47-L48 (Fig. 4B). We also observed a small peak for the product corresponding to L49-F50 cleavage. However, no peak corresponding to the cleavage between L48-L49 was observed. Further analysis led to the realization that with the natural Phe present at position 50 in WT CDH1, installing the designed L51F mutation generated 50F,51F CDH1, with two consecutive Phe at P1’ and P2’ relative to the WT ε-cleavage site. We previously showed that with 50F,51F mutants of APP^5^ and Notch,^31^ the ε-cleavage site shifts towards N-terminus by two amino acids, avoiding placement of either Phe in the P2’ position. Hence, the shift in the ε-cleavage site that we observed for L51F CDH1 is in fact consistent with our previous observations about 50F,51F mutants of APP and Notch as well as with the Phe specificity rule. However, the other observed CDH1 C-terminal product resulted from cleavage at L49-F50, placing the designed mutant L51F in the P2’ position. This apparent violation of the Phe specificity rule was investigated further through western blot analysis, to assess the effect of Phe mutation on total CDH1 C-terminal product formation.

### P2’ phenylalanine mutation reduces formation of N-terminal and C-terminal products

To further study the effect of Phe in substrate TMD on the efficiency of cleavage by γ-secretase, we performed immunoblotting analysis of C-terminal and N-terminal products followed by densitometry (Fig. 5). The C-terminal products were detected with anti-FLAG antibody and N-terminal products were detected using anti-HA antibody. The enzyme-substrate reactions were again performed, but for mutant and WT substrates in parallel. The reactions with only enzyme, only substrate, and enzyme plus substrate in the presence of inhibitor were also performed in parallel as controls.

**Figure 5:**
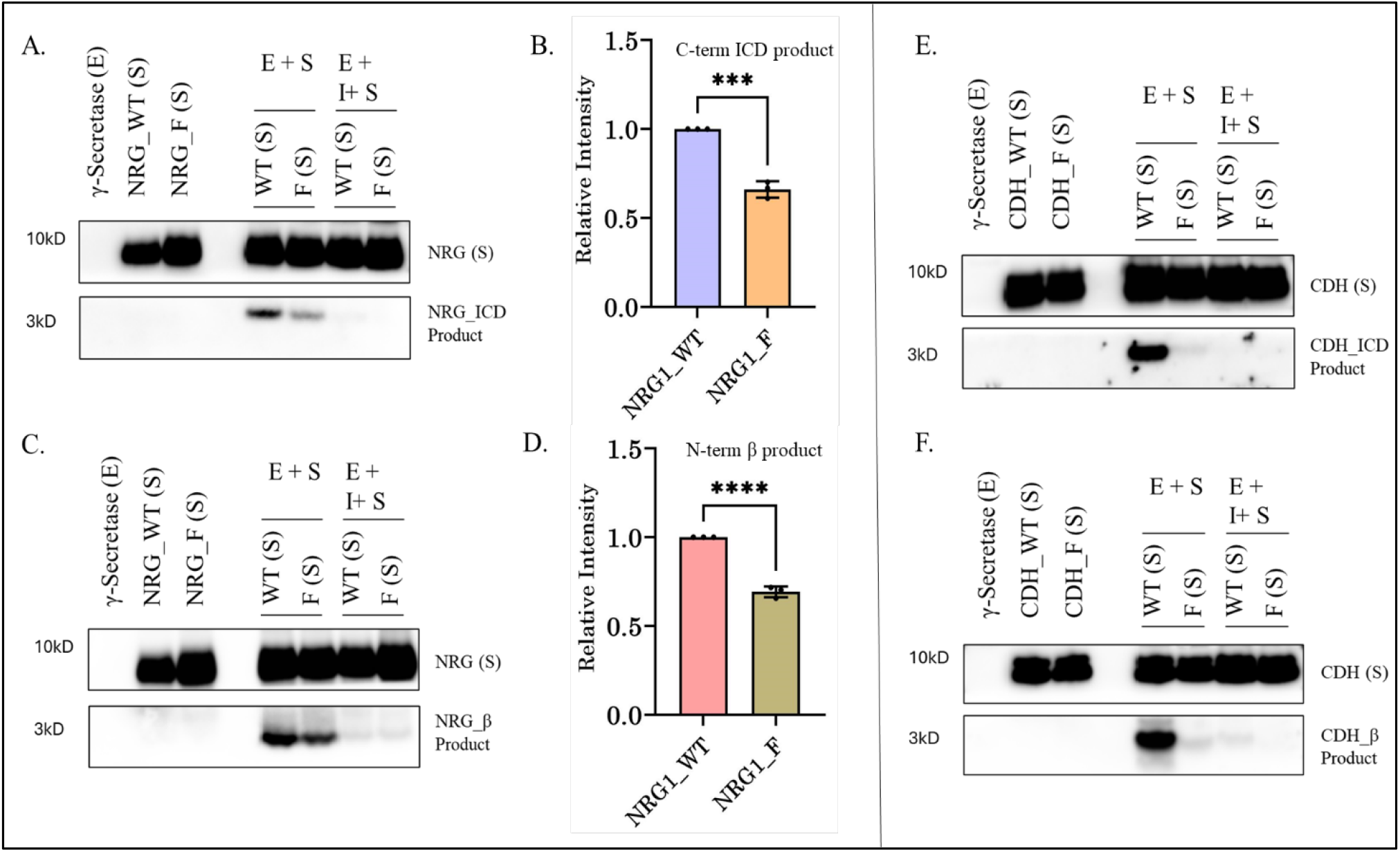
Immunoblots of γ-secretase cleavage products from phenylalanine mutants of NRG1 and CDH1. A. Immunoblot showing formation of C-terminal ICD-like products from NRG1 variants using anti-Flag antibody. The ICD-like product formation is reduced in the enzyme reaction with V51F mutant of NRG1 compared to WT. B. Densitometry of the ICD-like product band from WT and V51F NRG1 reactions. C. Immunoblot showing formation of N-terminal NRG1 β-like products using anti-HA antibody. NRG1 β product formation is reduced in the enzyme reaction with V51F mutant of NRG1 compared to WT. D. Densitometry of the NRG1β-like product band from WT and V51F NRG1 reactions. E. Immunoblot showing formation of the C-terminal ICD-like products from CDH1 variants using anti-Flag antibody. The ICD-like product formation is almost negligible in the enzyme reaction with L51F mutant of CDH1 compared to WT. F. Immunoblot showing formation of N-terminal CDH1 β-like products using anti-HA antibody. CDH1 β product formation is almost absent in the enzyme reaction with L51F mutant of CDH1 compared to WT. In all the blots, no product formation was observed in the control reactions of enzyme alone (E), substrate alone (S), or enzyme + substrate + inhibitor (E+S+I). N=3 Error bars= S.D. for each blot, and unpaired two-tailed student t-test was performed, where * *p* ≤ .05, ** *p* ≤ .01, *** *p* ≤ .001, **** *p* ≤ .0001. Densitometry was not performed on CDH1 blots as almost no product formation was seen from the mutant reaction. See Fig. S2 for full blots.

For V51F NRG1, C-terminal product formation was lowered to ~66% compared to WT (Fig. 5A-B); while N-terminal product was reduced to ~69% compared to WT (Fig. 5C-D). No product bands were observed in the control reactions in the C-terminal product blot (Fig. 5A), while very light bands appeared in the N-terminal product blots for substrate + enzyme reaction in presence of inhibitor (Fig. 5C), confirming γ-secretase-dependent product formation. In the case of L51F CDH1 (50F,51F CDH1), N-terminal and C-terminal product formation was all but absent, and the amounts produced were negligible and not conducive to quantification. Very light signals were observed after the long exposure of the blot that confirmed the formation of the products, although in a very low amount (Fig. 5E-F). This observation however is not surprising and consistent with our previous observation for γ-secretase cleavage of 50F,51F APP substrate.^5^ Despite the extremely low level of product formation, immunoprecipitation led to concentration of the C-terminal products, allowing for their detection and identification (Fig. 4B). No product bands were observed in the control reactions in the C-terminal blot (Fig. 5E), while for CDH-WT + enzyme reaction in presence of inhibitor, a very faint band was observed in the N-terminal product blot (Fig. 5F), confirming γ-secretase-dependent product formation.

## DISCUSSION

Although over 145 substrates for γ-secretase have been identified, their proteolytic processing is poorly understood except for the extensively studied APP followed by Notch1. We previously investigated the effects of TMD Phe residues in APP and Notch1 substrates on γ-secretase proteolysis.^5,31^ Here we extended this study for two other biologically important γ-secretase substrates: neuregulin1 (NRG1) and E-cadherin (CDH1). We found that WT NRG1 is cleaved at two different initial ε-sites, one the previously reported C49-V50,^19^ and a novel secondary cleavage between G46-I47. The reason for this second site is unknown. In contrast, ε-cleavage of WT CDH1 occurred only between L49-F50, instead of the previously reported L51-R52.^25^ This newfound ε-cleavage site for CDH1 is consistent with the Phe specificity rule, which avoids putting the natural F50 in the P2’ position for γ-secretase cleavage. Furthermore, this cleavage occurs within the TMD of the substrate, as commonly observed with other substrates and consistent with the intramembranous active site, containing two TMD catalytic aspartates. All the cleavages are summarized in Fig. 6.

**Figure 6:**
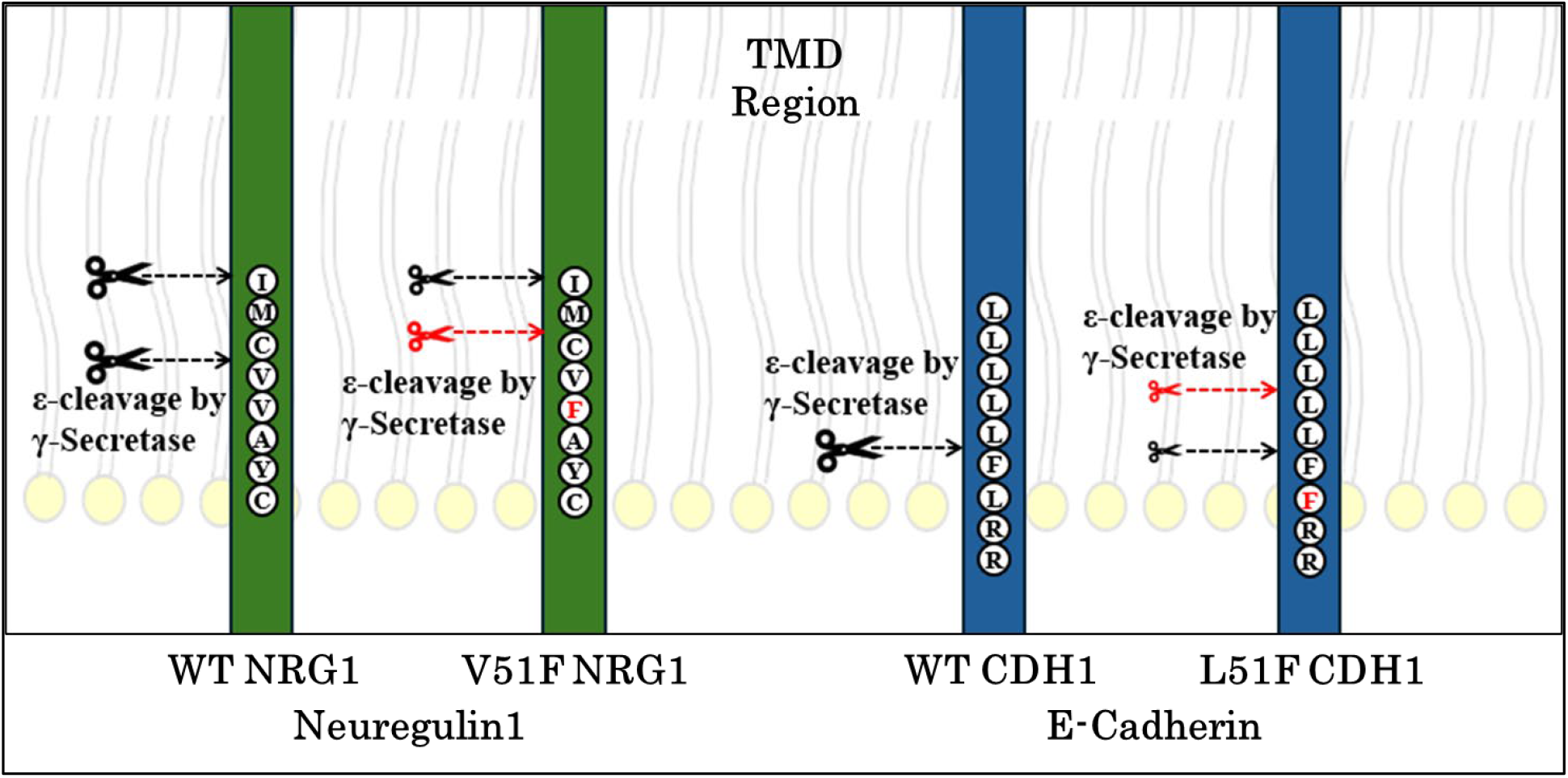
Summary of observed cleavages of WT and P2’ Phe mutants of NRG1 and CDH1 by γ-secretase. The red scissors indicate the shifted ε-cleavage site because of the P2’ Phe, which is also marked red in the figure, while small scissors indicate reduced proteolysis.

To determine if the Phe specificity rule for the P2’ position—applicable to APP and Notch1— holds true for NRG1 and CDH1, we mutated residues V51 in NRG1 and L51 in CDH1 to Phe. These residues mark the P2’ position relative to the ε-cleavage site observed here with the respective WT substrates (Table 1 and 2). For V51F NRG1, we observed the complete shift in the ε site by one amino acid in the N-terminal direction, i.e. between M48-C49. However, we still observed cleavage between G46-I47. No product corresponding to C49-V50 cleavage was detected. All these observations are consistent with the Phe P2’ specificity rule. For CDH1, given the natural Phe present at position 50, the L51F mutant generated 50F,51F CDH1; that is, with two adjacent Phe residues. Consistent with the Phe P2’ specificity rule, for L51F CDH1 (50F,51F) we observed a shift in the ε site by two amino acid in the N-terminal direction, i.e. between L47-L48, analogous to the cleavages reported previously for 50F,51F mutants of APP and Notch. No product corresponding to L48-L49 cleavage was observed. Surprisingly, we also observed a small peak of cleavage between L49-F50, placing Phe in P2’ position. However, as noted below, the total ε-cleavage of this double Phe substrate was negligible.

The immunoblots for C-terminal and N-terminal product analysis using anti-FLAG and anti-HA antibodies, respectively, showed reduction in product formation for the Phe-mutant substrates compared to WT with both NRG1 and CDH1. For V51F NRG1, both the C-terminal and N-terminal products were reduced to 65-70% compared to WT. In contrast, product formation for L51F CDH1 was too low to quantify either band intensities compared to WT. This dramatically reduced proteolysis of L51F CDH1 is likely due to the double Phe residues at the normal ε-cleavage site and is consistent with our previous observation made for the 50F,51F APP substrate mutant.^5^ Of note, knowledge of the correct ε-cleavage site for CDH1 and the complete blocking of CDH1 ε-cleavage by L51F enables generation of appropriate knock-in cells and mice to determine physiological consequences of specifically blocking CDH1 ICD production, among all γ-secretase substrates, and downstream signaling functions.

Taken together, all these data support the Phe specificity rule as general for γ-secretase substrates, that Phe is not tolerated at the P2’ position for cleavage by γ-secretase. Moreover, TMD Phe residues are also inhibitory, as this effect has now been observed with four γ-secretase substrates: APP, Notch1, NRG1 and CDH1. Whether the NRG1 and CDH1 substrates follow a processive tripeptide cleavage pattern is still unknown, as it is for Notch1. Answering this question will require detection of the small peptide co-products, as has been done to date only with APP substrate using LC-MS/MS.^4,41,42^

## ACCESSION CODES

UniProt Accession IDs for proteins used-NRG1:Q02297, CDH1: P12830, Presenilin-1: P49768, Aph1: Q96BI3, Nicastrin: Q92542, Pen2: Q9NZ42.

## ACKNOWLEDGEMENTS

We thank Dr. Eden Go (University of Kansas Synthetic Chemical Biology Core Laboratory) for technical support on mass spectrometry. This work was supported by Grant AG66986 from the National Institutes of Health to M.S.W. and Grant 2121063 from National Science Foundation to Yinglong Miao.

## AUTHOR CONTRIBUTIONS

S.R.M. performed all the experimental procedures. M.S.W. supervised the project. S.R.M. and M.S.W. analyzed and interpreted the data as well as wrote the manuscript. Both authors contributed to the final version of this manuscript.

## SUPPORTING INFORMATION (SI)

MALDI-TOF MS spectra for purified intact Phe mutants of NRG1 and CDH1, additional immunoblot images for substrates and their ICDs

## REFERENCES

1. Wolfe, M. S. (2019). Structure and Function of the γ-Secretase Complex. Biochemistry, 58(27), 2953–2966. 10.1021/acs.biochem.9b00401

2. Wolfe, M. S., Xia, W., Ostaszewski, B. L., Diehl, T. S., Kimberly, W. T., & Selkoe, D. J. (1999). Two transmembrane aspartates in presenilin-1 required for presenilin endoproteolysis and γ-secretase activity. Nature, 398(6727), 513–517. 10.1038/19077

3. Güner, G., & Lichtenthaler, S. F. (2020). The substrate repertoire of γ-secretase/presenilin. Seminars in Cell & Developmental Biology, 105, 27–42. 10.1016/j.semcdb.2020.05.019

4. Takami, M., Nagashima, Y., Sano, Y., Ishihara, S., Morishima-Kawashima, M., Funamoto, S., & Ihara, Y. (2009). γ-Secretase: Successive Tripeptide and Tetrapeptide Release from the Transmembrane Domain of β-Carboxyl Terminal Fragment. The Journal of Neuroscience, 29(41), 13042–13052. 10.1523/JNEUROSCI.2362-09.2009

5. Bolduc, D. M., Montagna, D. R., Seghers, M. C., Wolfe, M. S., & Selkoe, D. J. (2016). The amyloid-beta forming tripeptide cleavage mechanism of γ-secretase. eLife, 5, e17578. 10.7554/eLife.17578

6. Gazave, E., Lapébie, P., Richards, G. S., Brunet, F., Ereskovsky, A. V., Degnan, B. M., Borchiellini, C., Vervoort, M., & Renard, E. (2009). Origin and evolution of the Notch signalling pathway: An overview from eukaryotic genomes. BMC Evolutionary Biology, 9(1), 249. 10.1186/1471-2148-9-249

7. Meng, Y., Bo, Z., Feng, X., Yang, X., & Handford, P. A. (2024). The Notch Signaling Pathway: Mechanistic Insights in Health and Disease. Engineering, 34, 212–232. 10.1016/j.eng.2023.11.011

8. Siebel, C., & Lendahl, U. (2017). Notch Signaling in Development, Tissue Homeostasis, and Disease. Physiological Reviews, 97(4), 1235–1294. 10.1152/physrev.00005.2017

9. Shi, Q., Xue, C., Zeng, Y., Yuan, X., Chu, Q., Jiang, S., Wang, J., Zhang, Y., Zhu, D., & Li, L. (2024). Notch signaling pathway in cancer: From mechanistic insights to targeted therapies. Signal Transduction and Targeted Therapy, 9(1), 128. 10.1038/s41392-024-01828-x

10. Li, X., Yan, X., Wang, Y., Kaur, B., Han, H., & Yu, J. (2023). The Notch signaling pathway: A potential target for cancer immunotherapy. Journal of Hematology & Oncology, 16(1), 45. 10.1186/s13045-023-01439-z

11. Lemmens, K., Doggen, K., & De Keulenaer, G. W. (2007). Role of Neuregulin-1/ErbB Signaling in Cardiovascular Physiology and Disease: Implications for Therapy of Heart Failure. Circulation, 116(8), 954–960. 10.1161/CIRCULATIONAHA.107.690487

12. Falls, D. (2003). Neuregulins: Functions, forms, and signaling strategies. Experimental Cell Research, 284(1), 14–30. 10.1016/S0014-4827(02)00102-7

13. Longart, M., Calderón, C., González, M., Grela, M. E., & Martínez, J. C. (2022). Neuregulins: Subcellular localization, signaling pathways and their relationship with neuroplasticity and neurological diseases. Exploration of Neuroscience, 31–53. 10.37349/en.2022.00003

14. Dejaegere, T., Serneels, L., Schäfer, M. K., Van Biervliet, J., Horré, K., Depboylu, C., Alvarez-Fischer, D., Herreman, A., Willem, M., Haass, C., Höglinger, G. U., D’Hooge, R., & De Strooper, B. (2008). Deficiency of Aph1B/C-γ-secretase disturbs Nrg1 cleavage and sensorimotor gating that can be reversed with antipsychotic treatment. Proceedings of the National Academy of Sciences, 105(28), 9775–9780. 10.1073/pnas.0800507105

15. Zhu, X., Yu, G., Lv, Y., Yang, N., Zhao, Y., Li, F., Zhao, J., Chen, Z., Lai, Y., Chen, L., Wang, X., Xiao, J., Cai, Y., Feng, Y., Ding, J., Gao, W., Zhou, K., & Xu, H. (2024). Neuregulin-1, a member of the epidermal growth factor family, mitigates STING-mediated pyroptosis and necroptosis in ischaemic flaps. Burns & Trauma, 12, tkae035. 10.1093/burnst/tkae035

16. Wang, Y., Wei, J., Zhang, P., Zhang, X., Wang, Y., Chen, W., Zhao, Y., & Cui, X. (2022). Neuregulin-1, a potential therapeutic target for cardiac repair. Frontiers in Pharmacology, 13, 945206. 10.3389/fphar.2022.945206

17. Chambliss, C., Stiles, J. K., & Gee, B. E. (2023). Neuregulin-1 attenuates hemolysis- and ischemia induced-cerebrovascular inflammation associated with sickle cell disease. Journal of Stroke and Cerebrovascular Diseases, 32(2), 106912. 10.1016/j.jstrokecerebrovasdis.2022.106912

18. Yoo, J.-Y., Kim, H.-B., Lee, Y.-J., Kim, Y.-J., Yoo, S.-Y., Choi, Y., Lee, M.-J., Kim, I.-S., Baik, T.-K., Lee, J.-H., & Woo, R.-S. (2023). Neuregulin-1 reverses anxiety-like behavior and social behavior deficits induced by unilateral micro-injection of CoCl2 into the ventral hippocampus (vHPC). Neurobiology of Disease, 177, 105982. 10.1016/j.nbd.2022.105982

19. Fleck, D., Voss, M., Brankatschk, B., Giudici, C., Hampel, H., Schwenk, B., Edbauer, D., Fukumori, A., Steiner, H., Kremmer, E., Haug-Kröper, M., Rossner, M. J., Fluhrer, R., Willem, M., & Haass, C. (2016). Proteolytic Processing of Neuregulin 1 Type III by Three Intramembrane-cleaving Proteases. Journal of Biological Chemistry, 291(1), 318–333. 10.1074/jbc.M115.697995

20. Bao, J., Wolpowitz, D., Role, L. W., & Talmage, D. A. (2003). Back signaling by the Nrg-1 intracellular domain. The Journal of Cell Biology, 161(6), 1133–1141. 10.1083/jcb.200212085

21. Willem, M. (2016). Proteolytic processing of Neuregulin-1. Brain Research Bulletin, 126, 178–182. 10.1016/j.brainresbull.2016.07.003

22. Oda, H., & Takeichi, M. (2011). Structural and functional diversity of cadherin at the adherens junction. Journal of Cell Biology, 193(7), 1137–1146. 10.1083/jcb.201008173

23. Shapiro, L., & Weis, W. I. (2009). Structure and Biochemistry of Cadherins and Catenins. Cold Spring Harbor Perspectives in Biology, 1(3), a003053–a003053. 10.1101/cshperspect.a003053

24. Colás-Algora, N., & Millán, J. (2019). How many cadherins do human endothelial cells express? Cellular and Molecular Life Sciences, 76(7), 1299–1317. 10.1007/s00018-018-2991-9

25. Marambaud, P. (2002). A presenilin-1/gamma-secretase cleavage releases the E-cadherin intracellular domain and regulates disassembly of adherens junctions. The EMBO Journal, 21(8), 1948–1956. 10.1093/emboj/21.8.1948

26. Ferber, E. C., Kajita, M., Wadlow, A., Tobiansky, L., Niessen, C., Ariga, H., Daniel, J., & Fujita, Y. (2008). A Role for the Cleaved Cytoplasmic Domain of E-cadherin in the Nucleus. Journal of Biological Chemistry, 283(19), 12691–12700. 10.1074/jbc.M708887200

27. Wong, S. H. M., Fang, C. M., Chuah, L.-H., Leong, C. O., & Ngai, S. C. (2018). E-cadherin: Its dysregulation in carcinogenesis and clinical implications. Critical Reviews in Oncology/Hematology, 121, 11–22. 10.1016/j.critrevonc.2017.11.010

28. Yu, W., Yang, L., Li, T., & Zhang, Y. (2019). Cadherin Signaling in Cancer: Its Functions and Role as a Therapeutic Target. Frontiers in Oncology, 9, 989. 10.3389/fonc.2019.00989

29. Kaszak, I., Witkowska-Piłaszewicz, O., Niewiadomska, Z., Dworecka-Kaszak, B., Ngosa Toka, F., & Jurka, P. (2020). Role of Cadherins in Cancer—A Review. International Journal of Molecular Sciences, 21(20), 7624. 10.3390/ijms21207624

30. Wolfe, M. S. (2020). Substrate recognition and processing by γ-secretase. Biochimica et Biophysica Acta (BBA) - Biomembranes, 1862(1), 183016. 10.1016/j.bbamem.2019.07.004

31. Malvankar, S., & Wolfe, M. S. (2024). Effects of transmembrane phenylalanine residues on γ-secretase-mediated Notch-1 proteolysis. bioRxiv, 2024.12.11.628002. 10.1101/2024.12.11.628002

32. Haapasalo, A., & Kovacs, D. M. (2011). The Many Substrates of Presenilin/γ-Secretase. Journal of Alzheimer’s Disease, 25(1), 3–28. 10.3233/JAD-2011-101065

33. Zhou, R., Yang, G., Guo, X., Zhou, Q., Lei, J., & Shi, Y. (2019). Recognition of the amyloid precursor protein by human γ-secretase. Science, 363(6428), eaaw0930. 10.1126/science.aaw0930

34. Lu, P., Bai, X., Ma, D., Xie, T., Yan, C., Sun, L., Yang, G., Zhao, Y., Zhou, R., Scheres, S. H. W., & Shi, Y. (2014). Three-dimensional structure of human γ-secretase. Nature, 512(7513), 166–170. 10.1038/nature13567

35. Struhl, G., & Adachi, A. (2000). Requirements for Presenilin-Dependent Cleavage of Notch and Other Transmembrane Proteins. Molecular Cell, 6(3), 625–636. 10.1016/S1097-2765(00)00061-7

36. Bolduc, D. M., Montagna, D. R., Gu, Y., Selkoe, D. J., & Wolfe, M. S. (2016). Nicastrin functions to sterically hinder γ-secretase–substrate interactions driven by substrate transmembrane domain. Proceedings of the National Academy of Sciences, 113(5). 10.1073/pnas.1512952113

37. Philip, A. T., Devkota, S., Malvankar, S., Bhattarai, S., Meneely, K. M., Williams, T. D., & Wolfe, M. S. (2019). Designed Helical Peptides as Functional Probes for γ-Secretase. Biochemistry, 58(44), 4398–4407. 10.1021/acs.biochem.9b00639

38. Fraering, P. C., Ye, W., Strub, J.-M., Dolios, G., LaVoie, M. J., Ostaszewski, B. L., Van Dorsselaer, A., Wang, R., Selkoe, D. J., & Wolfe, M. S. (2004). Purification and Characterization of the Human γ-Secretase Complex. Biochemistry, 43(30), 9774–9789. 10.1021/bi0494976

39. Wong, G. T., Manfra, D., Poulet, F. M., Zhang, Q., Josien, H., Bara, T., Engstrom, L., Pinzon-Ortiz, M., Fine, J. S., Lee, H.-J. J., Zhang, L., Higgins, G. A., & Parker, E. M. (2004). Chronic Treatment with the γ-Secretase Inhibitor LY-411,575 Inhibits β-Amyloid Peptide Production and Alters Lymphopoiesis and Intestinal Cell Differentiation. Journal of Biological Chemistry, 279(13), 12876–12882. 10.1074/jbc.M311652200

40. Bolduc, D. M., Selkoe, D. J., & Wolfe, M. S. (2017). Enzymatic Assays for Studying Intramembrane Proteolysis. In Methods in Enzymology (Vol. 584, pp. 295–308). Elsevier. 10.1016/bs.mie.2016.10.026

41. Devkota, S., Williams, T. D., & Wolfe, M. S. (2021). Familial Alzheimer’s disease mutations in amyloid protein precursor alter proteolysis by γ-secretase to increase amyloid β-peptides of ≥45 residues. Journal of Biological Chemistry, 296, 100281. 10.1016/j.jbc.2021.100281

42. Devkota, S., Zhou, R., Nagarajan, V., Maesako, M., Do, H., Noorani, A., Overmeyer, C., Bhattarai, S., Douglas, J. T., Saraf, A., Miao, Y., Ackley, B. D., Shi, Y., & Wolfe, M. S. (2024). Familial Alzheimer mutations stabilize synaptotoxic γ-secretase-substrate complexes. Cell Reports, 43(2), 113761. 10.1016/j.celrep.2024.113761

