## Supplementary material for "γ-Secretase-mediated endoproteolysis of neuregulin-1 and E-cadherin": Fig. S1, Fig. S2

γ-Secretase-mediated endoproteolysis of neuregulin1

and E-cadherin

**Figure S1.** The MALDI-TOF MS for intact purified mutant proteins (substrates) are shown, confirming the protein expression and purification. The table shows the calculated and observed masses for A.NRG_F B.CDH_F.

| Substrates | Calculated mass | Observed mass |
|---|---|---|
| V51F NRG1 (NRG_F) | 8162.494 | 8163.495 |
| L51F CDH1 (CDH_F) | 7698.997 | 7699.842 |

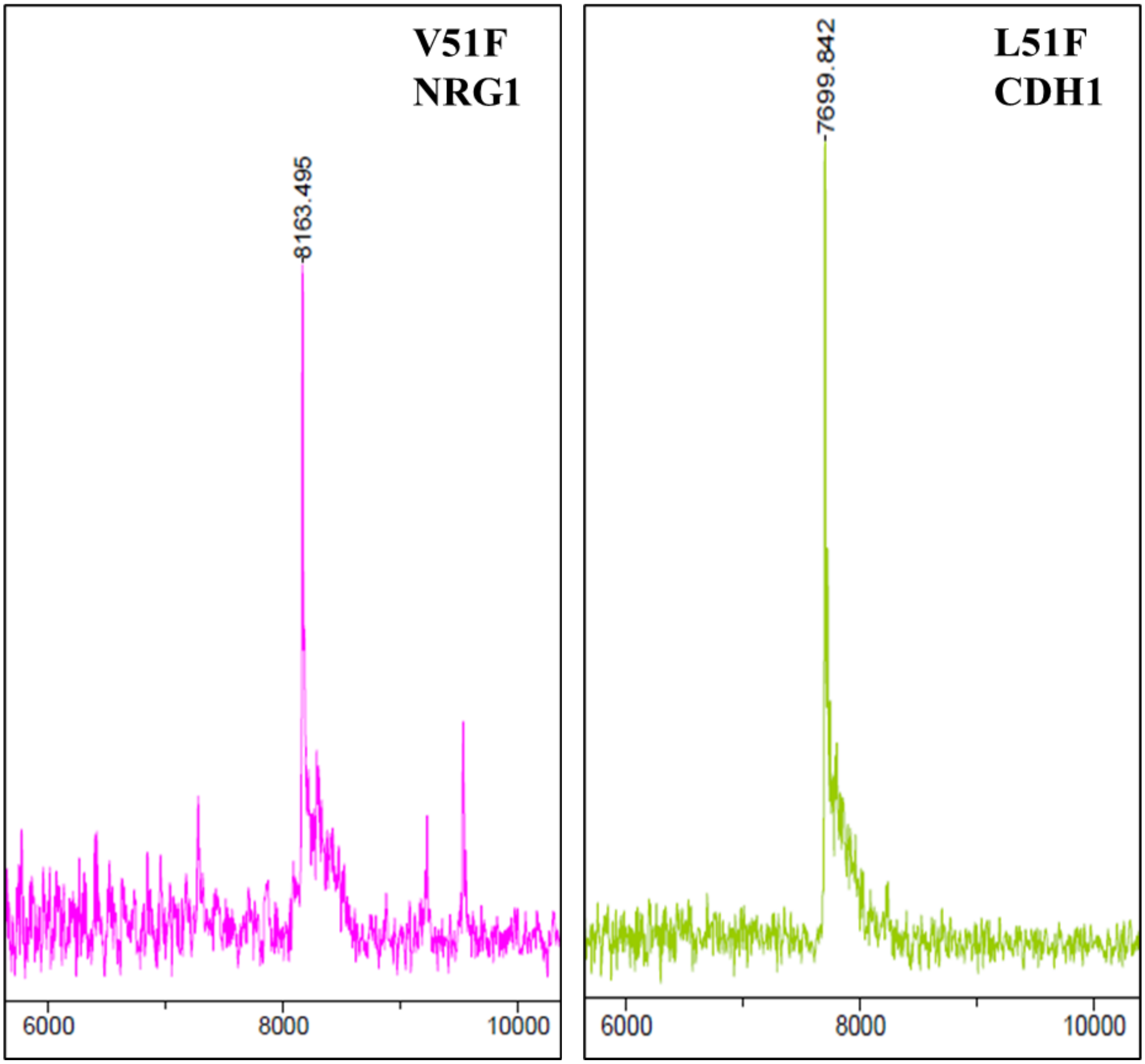

**Figure S2:** Additional immunoblots for analysis of γ-secretase processing of variants of neuregulin-1 (NRG1) and E-cadherin (CDH1) substrates. All blots in this figure were developed using anti-Flag antibody. As PEN-2 , a component of enzyme γ-secretase has Flag tag, the signal from PEN-2 is also visible.

**Section A:**

I. Full ICD Immunoblot for wild-type (WT) and Phe mutant of NRG1
    1. The low exposure blot shows uncleaved substrate bands for both NRG1 variants as well as PEN-2. However, the ICD from either of the NRG1 variants is not visible.
    2. The increase in exposure for the same blot increased the intensity of the uncleaved substrate and PEN-2 signals. Also, very light bands of the ICDs from WT and Phe mutant appeared.
    3. In the very high exposure of the same blot, the ICDs are visible but the quality of blot is diminished as uncleaved substrate signal is too intense, overpowering the blot.

II. Full ICD Immunoblot for wild-type and Phe mutant of CDH1
    1. The low exposure blot shows signal for intact CDH1 substrates (WT and Phe mutant) as well as for PEN-2. However, the ICD from either of the CDH1 variants is not visible.
    2. and 3. Higher exposures only intensified the signal from intact CDH1 substrates. However, even with higher exposures, the ICDs were not clearly visible for either CDH1 variant.

**Section B:** Full ICD immunoblot for WT NRG1 and CDH1 substrates

The full version of immunoblots shown in Fig. 3A and 3B of the manuscript are shown here. Specifically, the high exposure ICD blot in this section is added in the main manuscript as Fig. 3A and 3B.

The reactions and controls were run on the same gel for both WT substrates (NRG1 and CDH1). After transfer, the membrane was cut to separate bands of uncleaved substrates and ICD products and developed separately, to avoid the uncleaved substrate bands overwhelming the ICD product bands as seen in section A above. Furthermore, an enhancer solution was used for product bands to improve the signal of the bands, as mentioned in the Materials and Methods section. The low exposure and high exposure blots are shown. In addition to uncleaved substrate and ICD products, PEN-2 signal is also visible.

**Section A**

I. Full ICD Immunoblot for wild-type (WT) and Phe mutant of NRG1

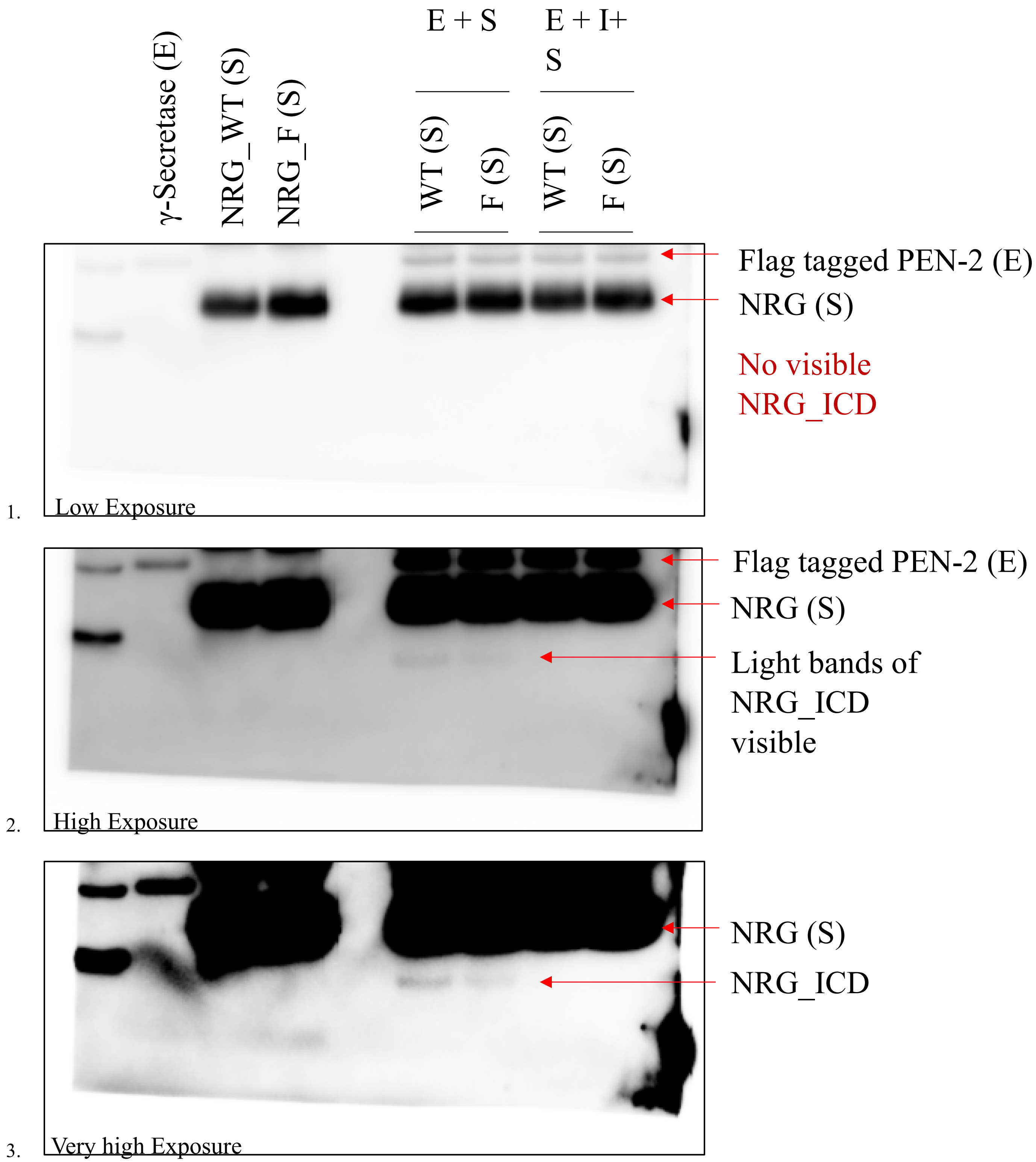

II. Full ICD Immunoblot for wild-type and Phe mutant of CDH1

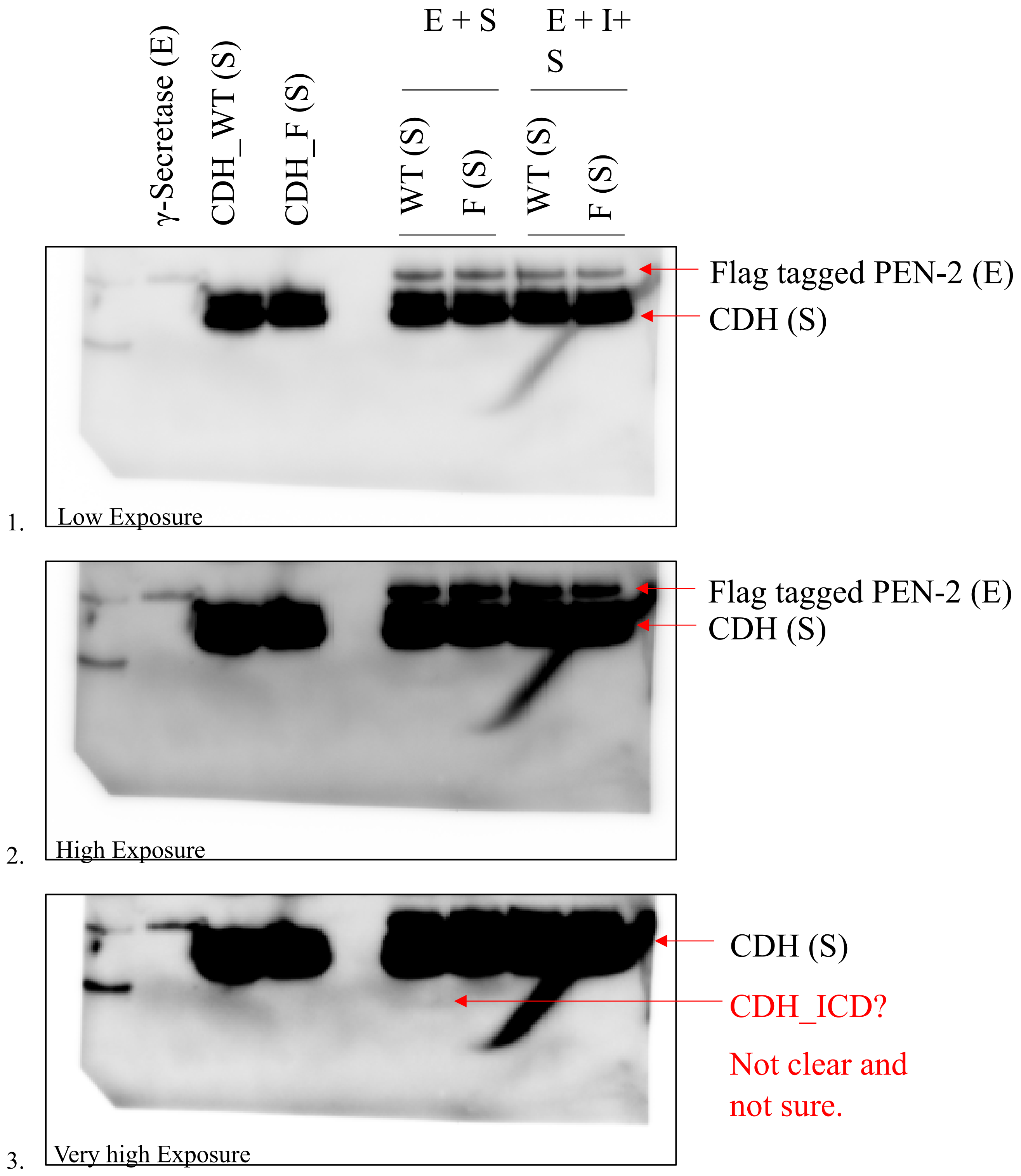

**Section B**

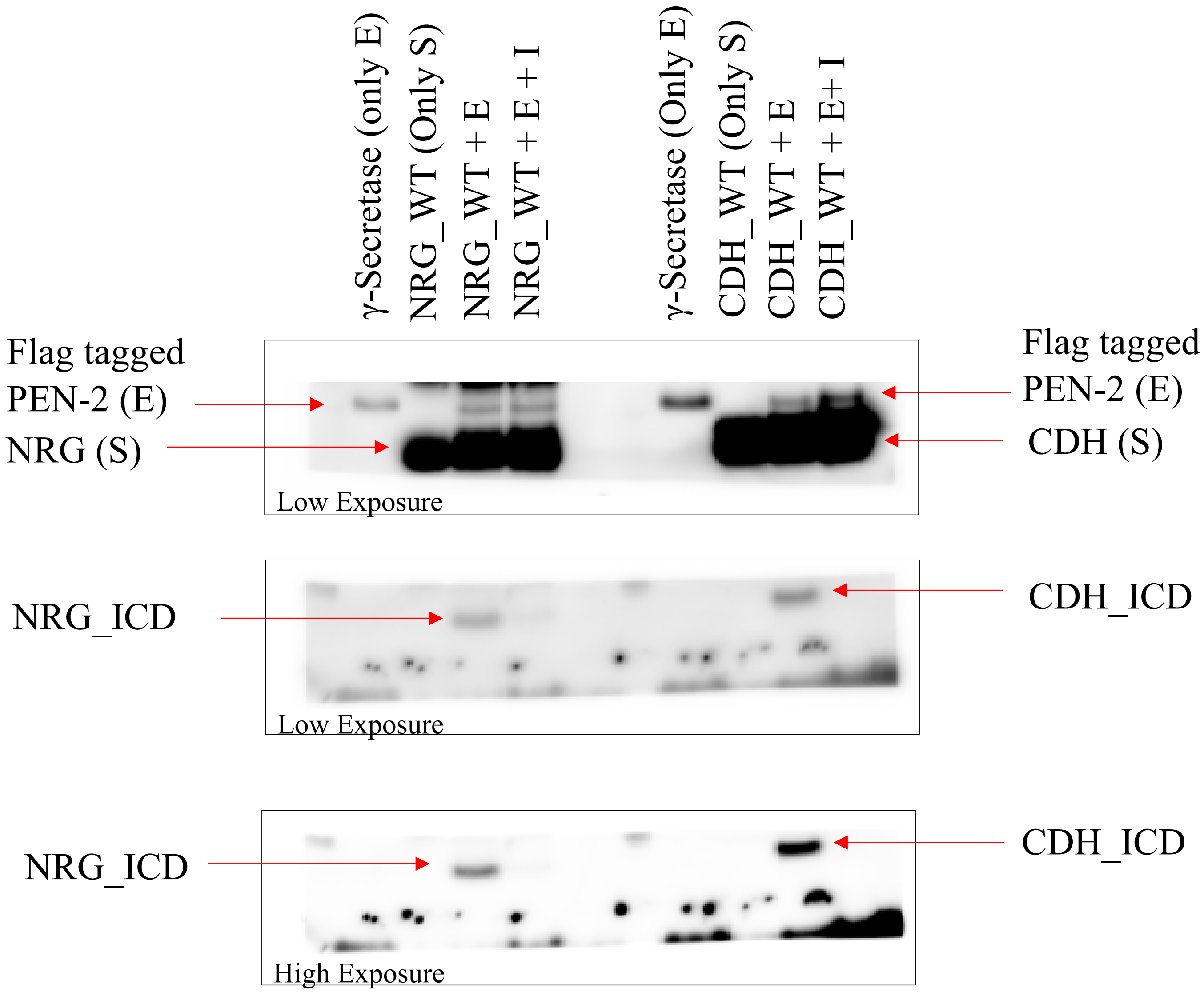
γ-Secretase (only E)
NRG_WT (Only S)
NRG_WT + E
NRG_WT + E + I
γ-Secretase (Only E)
CDH_WT (Only S)
CDH_WT + E
CDH_WT + E+ I
Flag tagged
PEN-2 (E)
NRG (S)
Flag tagged
PEN-2 (E)
CDH (S)
Low Exposure
NRG_ICD
CDH_ICD
Low Exposure
NRG_ICD
CDH_ICD
High Exposure
